# Management performance mapping and the value of information for regional prioritization of management interventions

**DOI:** 10.1101/380352

**Authors:** C. E. Buddenhagen, J. Andrade Piedra, G. A. Forbes, P. Kromann, I. Navarrete, S. Thomas-Sharma, Y. Xing, R. A. Choudhury, K. F. Andersen, E. Schulte-Geldermann, K. A. Garrett

## Abstract

Policymakers and donors often need to identify the locations and settings where technologies are most likely to have important effects, to increase the benefits from agricultural development or extension efforts. Higher quality information may help to target the high-payoff locations. The value of information (VOI) in this context is formalized by evaluating the results of decision making guided by a set of information compared to the results of acting without taking the information into account. We present a framework for management performance mapping that includes evaluating the VOI for decision making about geographic priorities in regional intervention strategies, in case studies of Andean and Kenyan potato seed systems. We illustrate use of Bayesian network models and recursive partitioning to characterize the relationship between seed health and yield responses and environmental and management predictors used in studies of seed degeneration. These analyses address the expected performance of an intervention based on geographic predictor variables. In the Andean example, positive selection of seed from asymptomatic plants was more effective at high altitudes in Ecuador. In the Kenyan example, there was the potential to target locations with higher technology adoption rates and with higher potato cropland connectivity, i.e., a likely more important role in regional epidemics. Targeting training to high performance areas would often provide more benefits than would random selection of target areas. We illustrate how assessing the VOI can help inform targeted development programs and support a culture of continuous improvement for interventions.

---

A central problem in applied spatial ecology is how to partition management efforts across landscapes. Interventions by governments or development organizations are often designed to increase regional crop yield, for example by improving disease management. International, governmental, and non-governmental organizations that seek to reduce poverty, enhance food security, and support ecosystem services, need strategies to geographically target interventions after identifying priorities using participatory approaches with stakeholders. We propose “management performance mapping” as a tool for translating experimental results to support identification of geographic priorities by policy makers and donors. Management performance mapping consists of scaling up models based on an often limited number of observations, to visualize how specific interventions are likely to perform at a regional scale (Altieri and Nicholls 2008; van Bussel et al. 2015; van Wart, Kersebaum, et al. 2013; Grassini et al. 2015). Management performance mapping can have a number of applications, such as providing a summary of recommendations for extension programs, or evaluating which type of management is most effective for a set of locations. In this paper, we focus on management performance mapping to inform targeting of interventions to support a management component known to be effective under some circumstances, where the goal is to identify the locations where it will be most effective. This approach may be particularly useful in low-income countries where smallholder farmers have fewer options, and there is interest in making a valuable new option available through a system intervention. Management performance mapping can be implemented to visualize the impact of proposed interventions, to improve decision-making and policymaking, as a component of adaptive management in development (Fig. 1).

**Figure 1.**
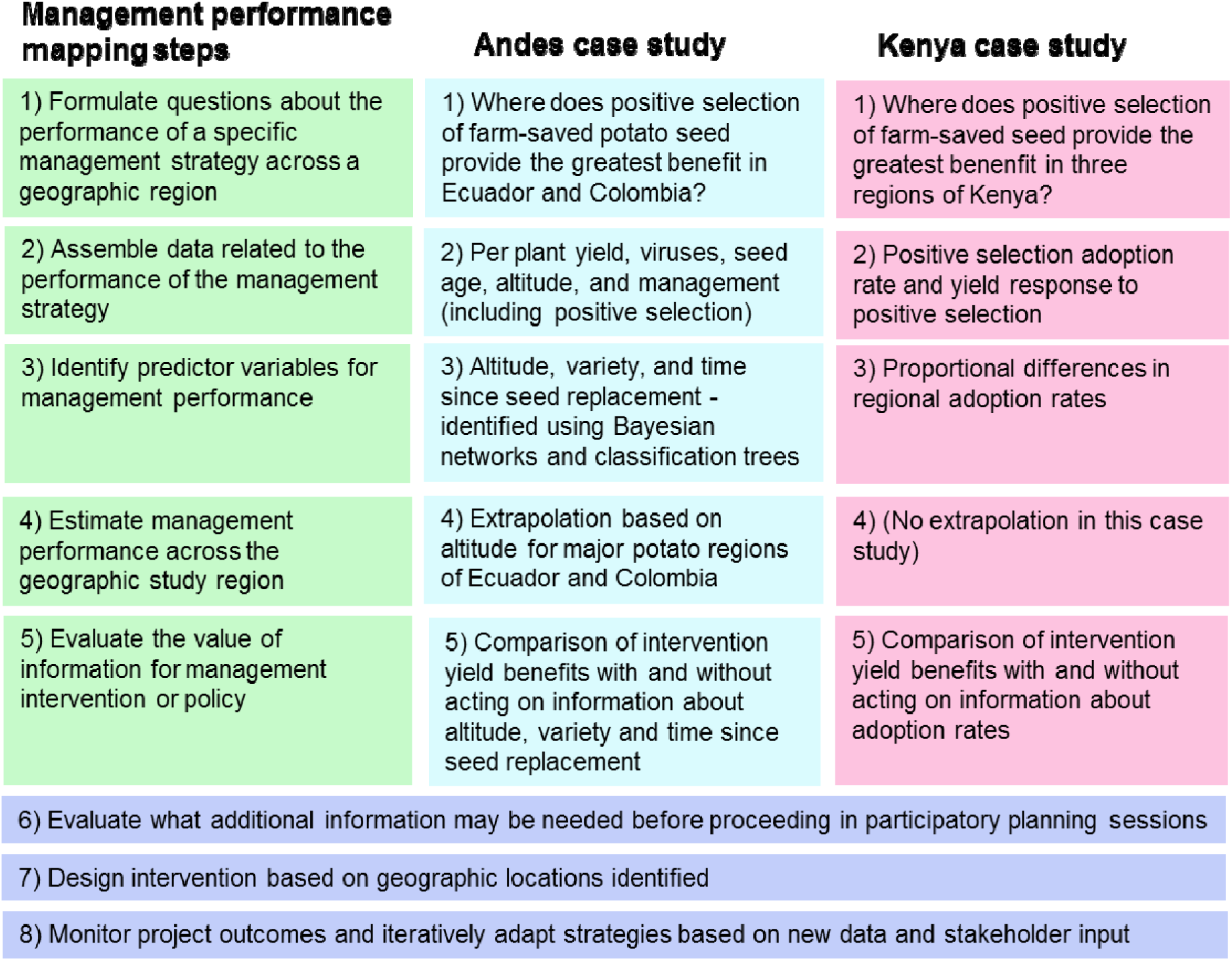
Steps in the management performance mapping pipeline. Selected development interventions should ideally take place in a culture of continuous improvement, based on ongoing monitoring and evaluation with stakeholders, and incorporating experimentation to facilitate adaptive management. Two case studies show how the steps may be implemented. Management performance mapping operates in this context by scaling up field, farm, and plot derived information to larger scale landscapes, regions or countries.

Digital or precision agriculture and species distribution models both address components of spatial prioritization and are thus related to management performance mapping. The question of how to optimize information use for decision making is addressed at the within-field scale in precision agriculture (Tittonell and Giller 2013), allowing well-resourced farmers, and potentially smallholder farmers (Cook et al. 2003), to collect and utilize spatially explicit data sets (in near real-time) about crop performance. Inputs such as fertilizer, pesticides, and irrigation are applied to areas of the field where they are most needed to maximize yields.

Species distribution models address the problem of optimal targeting indirectly, by providing information about where invasive (or endangered) species, including pathogens, are most likely to be found (Austin 2007; Hijmans and Graham 2006; Sheppard et al. 2014), often grappling with problems in statistical inference (Stolar and Nielsen 2015) also relevant to management performance mapping. Species distribution models are generally designed to draw inference beyond the regions where data were collected, by estimating species niche parameters based on maps of species occurrence or abundance throughout a species’ native and introduced range (Sutherst and Maywald 1985; Wang et al. 2017; Phillips et al. 2018; Bourdôt and Lamoureaux 2019). Management performance mapping for disease management can incorporate both information about which environments are conducive to pathogen and vector reproduction, and which environments are conducive to effective management.

The value of information (VOI) concept is useful for evaluating the benefits of basing strategies on management performance mapping. Assessing the VOI involves quantifying the expected benefit of reducing uncertainty (Canessa et al. 2015), as described further below. VOI analyses offer a means of both evaluating information and benefits, and assessing the role of uncertainty when comparing management options (Hirshleifer and Riley 1979; Macauley 2006; Canessa et al. 2015). VOI analyses compare outcomes from decision making with and without particular units of information, taking into account the stakes for making good or bad decisions, such as differences in yield or profit (Fig. 2). In studies of willingness-to-pay, such as farmer willingness-to-pay for technologies, the utility functions for technologies are closely related to the VOI (Breidert et al. 2006; Asante Bright Owusu et al. 2011; Hanemann 1991). Of course, decision-maker willingness-to-act based on information is necessary for information valuation to be meaningful. For example, overly confident decision-makers may not be influenced by new information, or they may not reflect on the uncertainty that is inherent in the information available. Many examples in the VOI literature focus on agriculture, such as the uncertainty risk distribution for farm yield (Hirshleifer and Riley 1979), the value of weather forecasting for farmers (Lave 1963), and risk assessment for crop futures (Danthine 1978). A related area of application of VOI concepts is in invasion biology more generally and in conservation biology, where decisions must also be made about where to prioritize efforts (Canessa et al. 2015; Johnson et al. 2017; Wilson 2015). VOI analyses have so far seen little application in plant pathology, crop epidemiology, or seed system development, where they have the potential to improve research prioritization and decision making.

**Figure 2.**
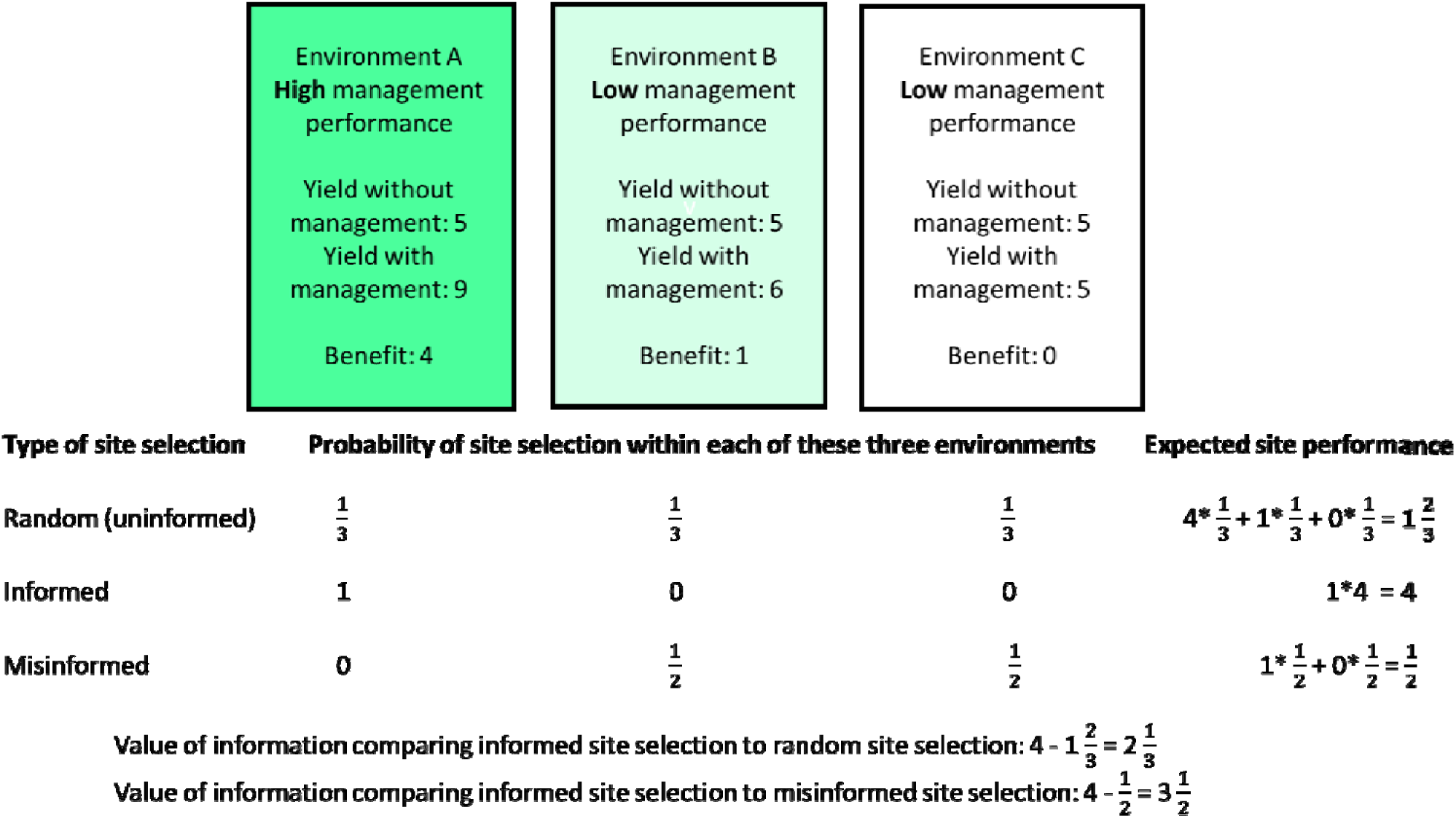
The of information (VOI) for data used to guide site selection for interventions can be evaluated as illustrated here for a hypothetical case. Suppose there are three types of environment, each equally common, and a measure of how well the management being evaluated performs in each environment: improvements in yield in environments A, B, and C of 4, 1, and 0 units, respectively. If sites are selected at random for intervention, without information about yield in the different environments, the average benefit from management is 1 2/3 units. If sites are selected considering the information about better management performance in environment A, and thus only environment A is targeted, then the average benefit from management is 4 units. If misinformation leads to the incorrect belief that management is more effective in environments B and C and these environments are equally targeted, then the average benefit from management is 1/2 unit. The VOI comparing informed site selection to random site selection is 2 1/3 units. The VOI comparing informed site selection to misinformed site selection is 3 1/2 units.

We present case studies of management performance mapping and the application of VOI analysis that focus on smallholder management of “seed degeneration” in agricultural systems. Seed degeneration is the reduction in yield or quality caused by an accumulation of pathogens (often viruses) and pests in planting material over successive cycles of propagation, where vegetatively-propagated crops deserve particular attention because of their higher risk of disease transmission (Thomas-Sharma et al. 2016). Establishing improved seed systems is challenging, especially in low-income countries, due in part to the many system components that must be integrated for seed system success (Jaffee and Srivastava 1994; McGuire and Sperling 2016; McQuaid et al. 2016; Sperling 2008; Gildemacher et al. 2009; Bentley and Vasques 1998; Almekinders et al. 2019). In informal seed systems in low-income countries, farmers typically use seed saved from the previous season for replanting, often leading to reduced yields, e.g., 5-50% reduction (Devaux et al. 2010a), especially when farmers are unfamiliar with approaches for selecting healthier seed from their fields with reduced pathogen risk. Despite the challenges (Almekinders et al. 2019), seed system improvement has great potential for improving regional agriculture, by providing healthier seed of better varieties, and has been a major focus of agricultural development efforts funded by many agencies (e.g., national plant protection agencies, The Bill and Melinda Gates Foundation, USAID, and FAO) (Jaffee and Srivastava 1994; McGuire and Sperling 2016; Almekinders et al. 1994).

Optimizing yield by reducing disease impacts, and improving seed quality, is a primary goal of many seed system interventions. Governments and institutions with a strong focus on science for development, such as CGIAR, work on a suite of factors linked to seed system health. Farmer training efforts focus on options for disease management and optimal decision-making. International development efforts for improved seed systems seek to increase farmer access to disease-free, disease-resistant, high-quality seed, to improve farmer practices, to implement “integrated seed health strategies” (Thomas-Sharma et al. 2016), and to implement realistic phytosanitary thresholds (Choudhury et al. 2017). Despite concerted efforts, many systems may revert to largely informal systems (sub-optimal seed sourced on-farm much of the time) after interventions. For example, 98% of potato seed sources in the Andes were reported as informal (Louwaars et al. 2013; Devaux et al. 2010a). Interventions are more likely to succeed if they are affordable, and help farmers to be profitable (McGuire and Sperling 2013; Sperling et al. 2013). As an example, positive selection is an on-farm management intervention that can provide large yield benefits, e.g., 28-55% increases (mean 32%) (Gildemacher et al. 2012, 2011; Schulte-Geldermann et al. 2012) and is often recommended as part of an integrated seed health strategy (Thomas-Sharma et al 2016). Under positive selection, farmers select healthy appearing plants and mark them for later harvesting of seed. Training farmers in the techniques of positive selection can be an effective component of an integrated seed health strategy, and we use positive selection as the example management in our case studies.

A challenge for management performance mapping – as for species distribution modelling, digital agriculture, and most analyses designed to draw inference about larger geographic areas – is to make the most of the available data while avoiding overinterpretation of results. Often data about agricultural management performance exist, or can be collected inside of existing intervention projects, but the data are collected at the scale of fields, farms or individual plant performance measures. Multiple factors influence plant productivity apart from management, generating uncertainty about the pay-off from management choices even where data are relatively abundant. We discuss considerations for use of limited data. The Andean case study addressed below illustrates both the challenge and potential value of management performance mapping. Greater vector activity is often assumed in lower elevations, suggesting that virus management in seed materials would be more important in these regions. Field observations in Ecuador, though based on a limited number of fields, suggest that the reverse is true for this case. We evaluate the VOI from management performance mapping to guide selection of intervention locations if this counterintuitive observation is indeed representative for the region.

Our objectives in this study are to (i) introduce and illustrate the concept of management performance mapping and associated methods, (ii) introduce the use of VOI analysis in this context, and (iii) illustrate the application of management performance mapping for potato seed degeneration management by positive selection of seed in the Andes and in Kenya. We also illustrate how analysis of likely management performance at individual sites can be combined with other geographic considerations, such as cropland connectivity as a proxy for the role of locations in epidemic spread for the region (Xing et al. 2020).

## METHODS

We describe the steps involved in producing management performance maps (Fig. 1), using the example of training farmers in positive selection to identify plants more likely to produce healthy seed. Then we illustrate management performance mapping for a seed degeneration data set from a potato seed study in Ecuador and a study of management adoption in Kenya (Gildemacher et al. 2012; Kromann et al. 2017). As a step in preparing the Andean management performance maps, we illustrate the application of Bayesian networks and recursive partitioning for assessing the influence of disease, environmental factors, and management on yield. We also evaluate the potential VOI for guiding the selection of locations in development interventions for potato seed health in Ecuador and Kenya based on the estimated effects, although we note that in these cases more data would be needed before proceeding to action in the field based on these analyses. We illustrate steps 1 through 5 of the management performance mapping pipeline (Fig. 1), while steps 6 through 8 would also be key to achieving outcomes in the field in an adaptive management approach (Shea et al. 2014). To illustrate the potential for combining management performance mapping (evaluated for each geographic pixel independently) with other types of spatial processes that may include the potential roles of locations in epidemic spread, we also provide an example of integration with a cropland connectivity analysis (Xing et al. 2020), described below.

### 1) Formulate questions about the performance of a specific management strategy across a geographic region

In these case studies, we evaluate the effects of positive selection of farm-saved seed potato for virus disease management. In general, the identification of management strategies for evaluation will likely be more successful if the process includes participatory input from stakeholders. In the Andean case study, our questions are: Where would training in positive selection likely produce the greatest benefit for yield in Ecuador and Colombia? And how does the variety grown and the time since seed replacement influence the benefit for yield? In the Kenyan case study, our question is: Where would training in positive selection likely produce the greatest benefit for yield, choosing among three regions of Kenya?

### 2) Assemble data related to the performance of the management strategy

We use two data sets as case studies. The first is from potato production in the Ecuadorian Andes, from a study designed for parameter estimation for a seed degeneration model. This study (Kromann et al. 2017) monitored seed degeneration in two potato cultivars, at three altitudes, and considered the use of on-farm seed management options. The two cultivars were INIAP-Fripapa and Superchola (perceived by farmers to be susceptible and resistant to degeneration, respectively). The field trials were carried out during three cycles of planting at three sites representing three altitudes (<2700 masl, 3000 masl and > 3400 masl, where the site <2700 masl was moved during the course of the experiment). Twelve 49 m^2^ plots were planted each year, two plots of each cultivar at each altitude/site. In each whole plot, three types of seed management were carried out in subplots: positive selection, roguing and random selection. The response variables included (1) virus incidence (Potato virus X (PVX), Potato virus Y (PVY), Potato virus S (PVS), Potato leaf roll virus (PLRV), Andean potato latent virus (APLV), and Andean potato mottle virus (APMoV)) in plants at emergence, flowering and in tubers, evaluated using DAS-ELISA, (2) incidence and severity of pest damage and diseases in tubers, and (3) tuber yield.

This Ecuadorian study was designed for parameter estimation for a seed degeneration model (Thomas-Sharma et al. 2017). The main components of this model relate to seed health (virus incidence, time/seasons since certified seed was last obtained), cultivar, environmental factors (weather), management (seed propagation and selection) and yield data for samples of individual potato plants (Kromann et al. 2017). A single site represented each altitude in this data set, so variability within a scenario can only be evaluated at the individual plant level. Lack of replication at the field level is a limitation for management performance mapping, because an analysis intended for providing recommendations for project implementation would be stronger if multiple farms per altitude provided estimates of farm-to-farm variation in management performance within an altitude range. We focus on yield data as the response in the management performance mapping example, with potential predictors being farm altitude (across three altitudes), seasons since certified seed was obtained, and the management performance of positive selection compared to roguing or random seed selection as management strategies. Climate variables – precipitation, humidity and temperature, from the WorldClim data base (Fick and Hijmans 2017) – were also evaluated as potential predictors, but were not effective predictors of either disease incidence or yield, probably at least in part because only three fields per year were evaluated (data and analysis not shown).

The second data set was published data about seed health management, and positive selection training and adoption rates in three counties in Kenya (Gildemacher et al. 2012). We used this data to illustrate integrating information about the likelihood that farmers in a region will adopt a technology (Gildemacher et al. 2012), another key component of intervention success.

### 3) Identify predictor variables for management performance

There are many potential predictor variables for performance indicators (Thomas-Sharma et al., in preparation) and a wide range of methods can be used to identify important predictors, including regression analysis, generalized linear models, and generalized additive models. We illustrate two types of machine learning algorithms – classification and regression trees, and Bayesian network analysis – to evaluate potential predictors, focusing on yield as the response used as a management performance indicator. These two methods were used to identify predictor variables for the effect of positive selection on yield for the Kromann et al. (2017) dataset. Simpler approaches to identifying key predictors may also often prove useful in application of management performance mapping.

#### Classification and regression trees

Classification and regression trees have been applied in agricultural systems for land and soil classification, climate change impact assessment, risk assessment, and evaluation of toxin levels and disease conduciveness in plants (Langemeier et al. 2016; Novak and LaDue 1999; Etter et al. 2006; Caley and Kuhnert 2006; Paul and Munkvold 2004; Tittonell and Giller 2013). The strength of the recursive partitioning method lies in its ability to deal with non-linearity in data and to depict and support interpretation of the outputs in a decision-tree format. A limitation of this method is that it may perform relatively poorly with continuous variables or large numbers of unordered variables. We illustrate use of the rpart package in R in the following two examples using the Kromann et al (2017) data set.

#### Effect of seed selection, time since seed renewal, and altitude on yield, evaluated with recursive partitioning (Andes)

We evaluated yield as the response variable, with predictors being the use of positive selection (as opposed to roguing or random seed selection), the time since seed renewal through purchase of certified seed (either three seasons or less than three seasons), and the effect of altitude (across three altitudes). Because altitude is available as a potential geographic predictor variable for the region, it is a candidate for extrapolating analysis of the performance of positive selection to a wider area in Step 4 (Fig. 1).

#### Effect of potato cultivar and its interactions on yield, evaluated with recursive partitioning (Andes)

In a previous study of a grower cooperative in Tungurahua, Ecuador, Superchola (one of the most important potato varieties in Ecuador) and INIAP-Fripapa were sold and grown at a ratio of approximately 2:1 by volume (Buddenhagen et al. 2017). Our analysis of the Kromann et al. dataset (2017) also focuses on these two varieties. We evaluate the effects of cultivar, altitude, and management by estimating mean per-plant yields across treatment combinations, and by using recursive partitioning in rpart.

#### Bayesian networks

Bayesian networks (Therneau et al. 2010) have been applied in natural resource management systems for applications such as vegetation classification, optimal decision making, disease management, adaptive management of wildlife habitat, and expert elicitation (Geenen and Van Der Gaag 2005; Aguilera et al. 2011; Kristensen and Rasmussen 2002; Perez-Ariza et al. 2012; Howes et al. 2010). A Bayesian network is a directed, acyclic graph whose nodes represent predictor variables and links represent dependencies. The relationships between variables are quantified in conditional probability tables, where the set of all tables together represents the full joint distribution. Important strengths of the Bayesian network method include its ability to infer probabilistic relationships among many variables simultaneously. The network structure can be set manually by the user or learned from the data using a variety of algorithms. In the case of exact estimation algorithms, it is possible to set values for any combination of nodes and produce new posterior probabilities for each variable in the network. A limitation of this method is the cost of some of the most advanced Bayesian network software. In addition, combinations of continuous and categorical data can be problematic for some commonly-used Bayesian network algorithms (Aguilera et al. 2011). Tools available for Bayesian network analysis include BI-CAMML, Hugin and Netica (Aguilera et al. 2011). R packages include bnlearn, gRain and pcalg (Nagarajan et al. 2013). We selected Netica for this illustration because it is relatively affordable, the algorithms it uses allow for immediate updating of conditional probabilities based on selected levels for variables, it has a powerful graphical interface, and it is widely used in ecological and environmental analyses (Aguilera et al. 2011).

#### Effect of positive selection on yield, evaluated in Bayesian networks (Andes)

The benefit of positive selection (in the third cropping cycle after certified seed purchase) was evaluated in a Bayesian network in Netica. Netica’s Tree-Augmented Naive Bayes (*TAN*) classifier algorithm was used to estimate the conditional probability tables and the network structure. From the conditional probability tables we estimated yields above (7.7 t/ha) and below (3.2 t/ha) the threshold altitude identified in the analysis: 2895 m.a.s.l.

#### A simple analysis of regional differences in adoption of training recommendations (Kenya)

In this case study, we evaluated regional differences in farmers’ adoption of positive selection after training (Table 1), reported by Gildemacher et al. (2012) as follows for three Kenyan counties: Nakuru 46%, Nyandarua 19%, and Narok 18%.

**Table 1.** Regional adoption rates after positive seed selection training in Kenya and the expected benefit of training given the adoption rate (Gildemacher et al. 2011, 2012). The average benefit is that expected under random allocation of training effort to the regions without regard to adoption rates.

| County | Observed adoption rate | Per household benefit \$USD | Expected benefit of training in year 1 |
| --- | --- | --- | --- |
| Nakuru | 0.46 | 156 | 72 |
| Nyandarua | 0.19 | 156 | 30 |
| Narok | 0.18 | 156 | 28 |
| Average Benefit: |  |  | 44 |

### 4) Estimate management performance across the geographic study region

For positive selection of on-farm seed in the Andes, we selected for analysis and extrapolation a major potato growing region stretching from southern Ecuador to southern Colombia. Using potato production geographic data layers from SPAM 2005 v3.2 Global Data (IFPRI and IIASA 2016) we focused on pixels with >200 ha potato production per pixel (where a pixel represents 5-arc minutes, approximately 10,000 ha). Here 51% of potato production is above 2895 m (the altitude threshold identified in the analyses above) based on SPAM estimates (You et al. 2012). The resulting management performance map will indicate that these regions would be priorities for targeting training in positive selection if decisions are based solely on this analysis of the data from Kromann et al. (2017).

For positive selection of on-farm seed in Kenya, rather than extrapolating the estimates of management performance for positive selection, we simply compare the relative performance of the counties (Table 1). The resulting management performance map will indicate prioritization among these counties if decisions for targeting positive selection training are based solely on the data from Gildemacher et al. (2009).

### 5) Evaluate the value of information for management intervention or policy

We assessed the value of information for decisions about where to invest management interventions, for a scenario where the estimates from Kromann et al. (2017) do correctly represent the region. For the purposes of this illustration, we considered cases where decision makers either have or do not have information about the geographic differences in management performance (Fig. 2). In the absence of information, they might select any location for management with equal probability. An estimate of the value of information would be the difference in the benefit of investment for locations selected based on the information (“informed location selection”), and the benefit for locations selected randomly (“uninformed location selection”). For example, informed site selection might direct site selection to farms above or below the altitude threshold identified in analysis (e.g., 2895 m.a.s.l. identified in Bayesian network analysis), depending on whether higher or lower elevations provide greater benefits. In the case where decision makers have a prior belief that is not supported by the data, and it is in fact an incorrect belief, the value of information would be the difference between investment outcomes based on the misconception (“misinformed location selection”) and outcomes based on informed investments. For example, there could be a prior belief that a particular pathogen will be more prevalent at lower elevations, due to a higher abundance of vectors, resulting in a prior belief that positive selection would be more important at lower elevations. We evaluated uninformed, informed, and misinformed management choices related to spatially distributed differences in yield, disease, cultivar and the rates with which best practices are adopted.

#### VOI for positive selection targeting in the Andes

Comparison of yield improvements due to positive selection training – with and without the information from Kromann et al. (2017) has as a first step determining how common each trait combination is in the landscape being considered. Then the probability of randomly including a particular trait combination can be estimated. The proportion of Ecuadorian farmers using certified seed was previously reported at 2% (Devaux et al. 2010b), and many farmers lack access to certified seed, though for some organized farming groups the proportion using certified or quality-declared seed can be as high as 46% (Buddenhagen et al. 2017). We take the frequency of farms in this landscape being planted with certified seed (“new seed”) at any given time as being approximately 2% (so that a farm drawn at random has probability p = 0.02 of being planted with certified seed, although this is an approximation because it is generally the wealthier farmers, government programs, or non-governmental organizations who acquire certified seed). Farm altitude, based on the geographic analysis described above for higher density potato regions, is above the altitude threshold identified in recursive partitioning approximately 51% of the time. For simplicity, we treat the potato cultivar planted as 33% INIAP-Fripapa and 66% Superchola, based on estimates for the province of Tungurahua from Buddenhagen et al. (2017).

#### VOI for targeting positive selection in Kenya

The average benefit of positive selection was reported by Gildemacher et al. (2012) as 3.4 tons per ha (∼$350 per ha). This translated to a per-household benefit of $156 per season for a farm of average size for the region. Meanwhile, the cost of training was $38 per farmer. In this case, the expected first-year benefit was $44 per household when training occurred in a randomly selected region (without regard to adoption rate) (Table 1). We compare this outcome to the outcome using information about frequencies of adoption.

### Integration with another criterion for selecting priority locations: cropland connectivity (Ecuador and Colombia)

The data layer of estimated management performance is one important factor for deciding where to prioritize management efforts. The management performance map developed up to this stage is generated pointwise, in that it treats each location (point) as independent from other locations. However, some locations will have more important roles in epidemics than others, due to factors such as the location’s position in spatial epidemic networks. Thus, targeting some locations will have more important effects to slow regional epidemics, for seed degeneration pathogens such as viruses that tend to be spread from one field to another. We also evaluated the layer of management performance estimates for positive selection integrated with a data layer of the potato “cropland connectivity risk index”, a measure of the likely importance of locations for spatial movement through potato growing areas (Xing et al. 2020; Margosian et al. 2009), as described below.

The potato cropland connectivity analysis was based on the potato crop harvested area data from SPAM 2005 v3.2 Global Data (IFPRI and IIASA 2016). This data has pixel resolution 5-arc min, and those cells with harvested area greater than 200 ha were included in the cropland connectivity risk analysis (Xing et al., 2020). As described in more detail in Xing et al. (2020), the distance between pairs of cells was evaluated in a sensitivity analysis for both inverse power-law models (parameters 0.5, 1, and 1.5) and negative exponential models (parameters 0.05, 0.1, 0.2, 0.3, and 1). Three network link thresholds (0.001, 0.0001, 0.00001) were applied separately to each adjacency matrix to represent three different scenarios in the network analysis in a sensitivity analysis. A cropland connectivity risk index (CCRI) was calculated as the scaled weighted sum of betweenness centrality, node strength, the sum of nearest neighbours’ node degrees, and eigenvector centrality, as in Xing et al. (2020). For each realization in the sensitivity analysis, the mean CCRI was evaluated across the 24 parameter combinations. This mean CCRI was then mapped in combination with the map of management performance estimates, to identify locations important both for the CCRI (indicating a potentially important epidemic role) and for management benefits from positive selection.

## RESULTS

### 3) Identifying predictor variables for management performance

#### Positive selection and yield for Andean potato

*In the recursive partitioning analysis*, higher per plant yields were generally obtained from INIAP-Fripapa (compared to Superchola) in the first two years after the certified seed was purchased, the highest yields being obtained for altitudes over 3278 m (Fig. 3). The highest yields for Superchola were found above 2895 m altitude. If a farmer can afford to replace seed more frequently, and the farm is over 3200 m, INIAP-Fripapa yielded higher than Superchola (and their value in the market was comparable in 2016 – 0.29 and 0.33 USD per kg, respectively).

**Figure 3.**
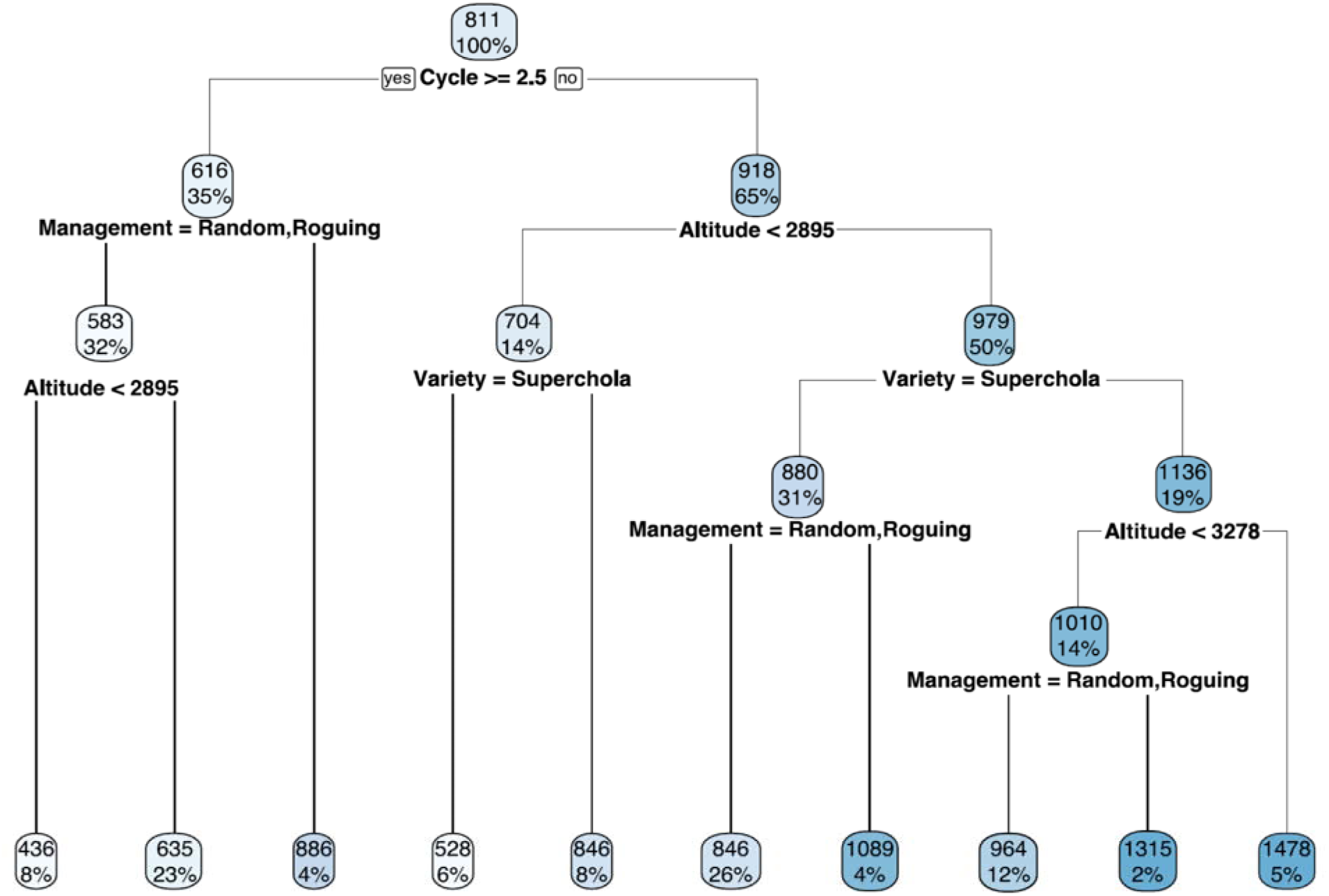
Recursive partitioning results in a decision tree format, with per plant yield (g) of potato in Ecuador as the response variable, based on the Kromann et al. (2017) dataset. Branches to the left are results when the logical statements at the nodes are true, and branches to the right are results when the logical statements are false. The upper numbers in the boxes are the mean yields for that condition, and the percent values are the proportion of the data for which the condition applies. For “Cycle >= 2.5”, “yes” indicates that the time since seed replacement with certified seed was greater than 2 years, while ‘no’ indicates that it was 2 or fewer years. For “Management = Random, Roguing”, “yes” indicates that either roguing or random seed selection was implemented, while ‘no’ to that option indicates that positive selection was implemented. For “Variety = Superchola”, “no” indicates that the variety was INIAP-Fripapa. (Darker colors indicate a higher number of ‘no’ answers for that condition compared to other conditions.)

Plants three years post-certified seed purchase yielded 33% less per plant. The benefits of positive selection allow yields to approach the average yields for recently purchased certified seed. There were no differences with respect to cultivar three years after certified seed purchase, suggesting positive selection was equally valuable in both varieties for seed that had gone through more than two planting cycles.

*The Bayesian network analysis* indicated that high yielding plants were found more commonly in plots where first generation certified seed was used, at higher altitudes, for the INIAP-Fripapa cultivar, and where there was a lower minimum temperature and higher rainfall six months after planting, as well as low levels of PVX, PLRV, and PVY (Fig. 4). Positive selection was less likely to be the management implemented if the yield was low. All the viruses except PVY, and PLRV for low yield plants, were more likely to be absent (frequency = 0) than present. Each virus was relatively more likely to be absent if a plant was in the high yield category compared to plants in the low yield category (Fig. 4). The uncertainty was high compared to the observed values, indicating that another cycle of data collection would be needed before implementing project plans based on this data.

**Figure 4.**
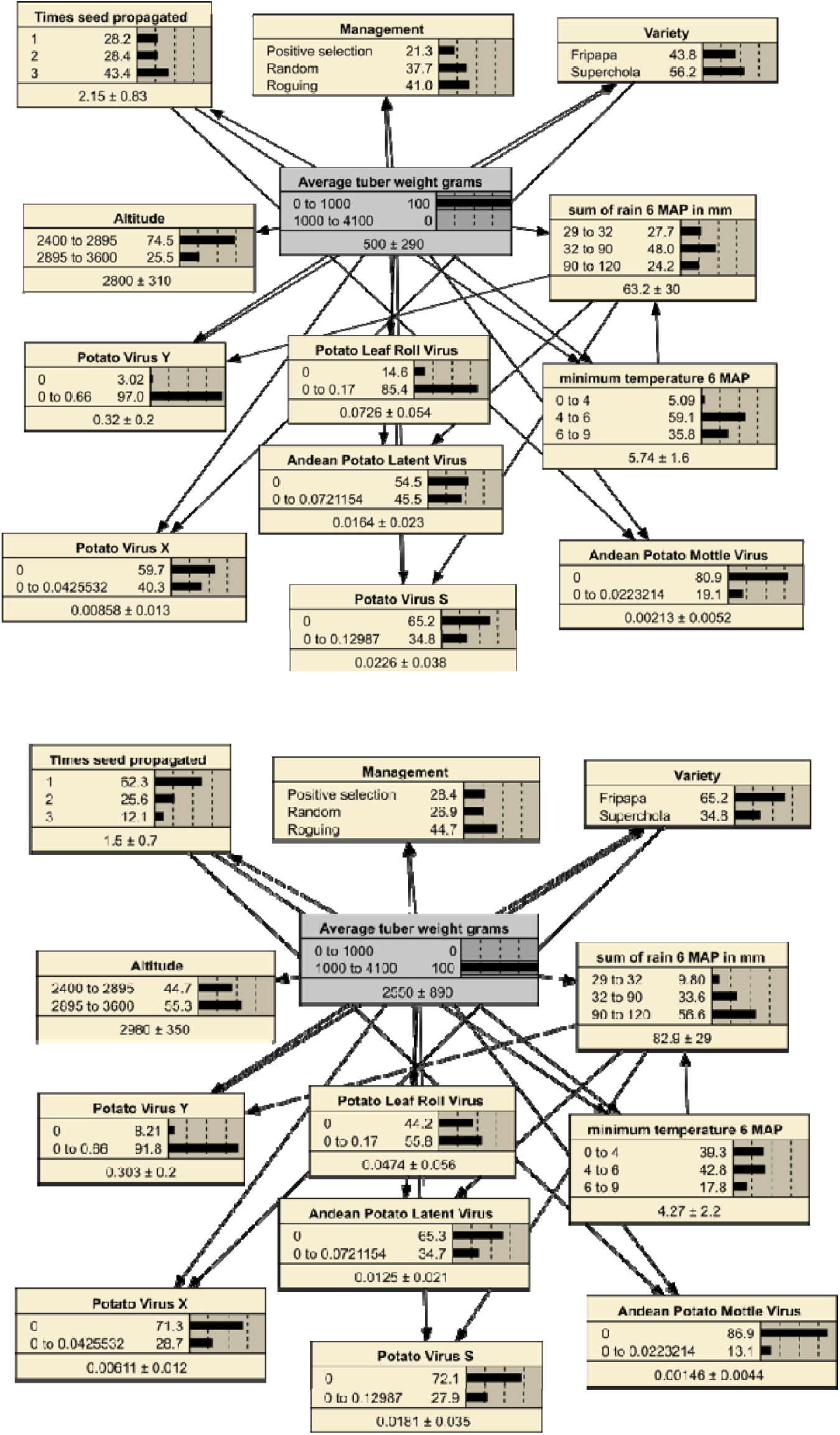
Ecuadorian potato yield and the factors associated with yield from a Bayesian network analysis carried in Netica using the Kromann et al. (2017) dataset. The two networks indicate the frequency distribution of a set of twelve potential predictor variables for plants with low yield (top) and high yield (bottom). The lower text for each node gives the estimated mean and uncertainty. Positive selection was less likely to be the management implemented for cases where the yield was low (top network). All the viruses except *Potato virus Y*, and *Potato leaf roll virus* for low yield plants, were more likely to be absent (frequency = 0) than present. Each virus was relatively more likely to be absent if a plant had high yield (bottom) than if it had low yield.

#### Adoption of positive selection for Kenyan potato

This analysis was based on the probability of adoption of positive selection, where higher adoption rates resulted in a higher payoff for intervention investment. Adoption rates were 46, 19 and 18% in three counties (Table 1) (Gildemacher et al. 2012, 2011). Thus, based on this measure alone, selection of the county with 46% adoption rate would approximately double the benefits obtained from a training intervention focused only where there was 18% or 19% adoption.

### 4) Estimating management performance across the geographic study region, and integrating with cropland connectivity estimates

#### Applying models to a map of the relevant region and integrating data layers for Andean potato

The mapped estimates of the management performance of positive selection for Andean potato yield (based on altitude) and the locations where potato cropland connectivity risk was highest based on the CCRI (Fig. 5) were combined to identify locations both (a) independently likely to have the best management outcomes, and (b) likely important for regional management of disease spread (high CCRI). Locations that meet both criteria were observed near the border of Ecuador and Colombia, and near Ambato and Riobamba in Ecuador (Fig. 5).

**Figure 5.**
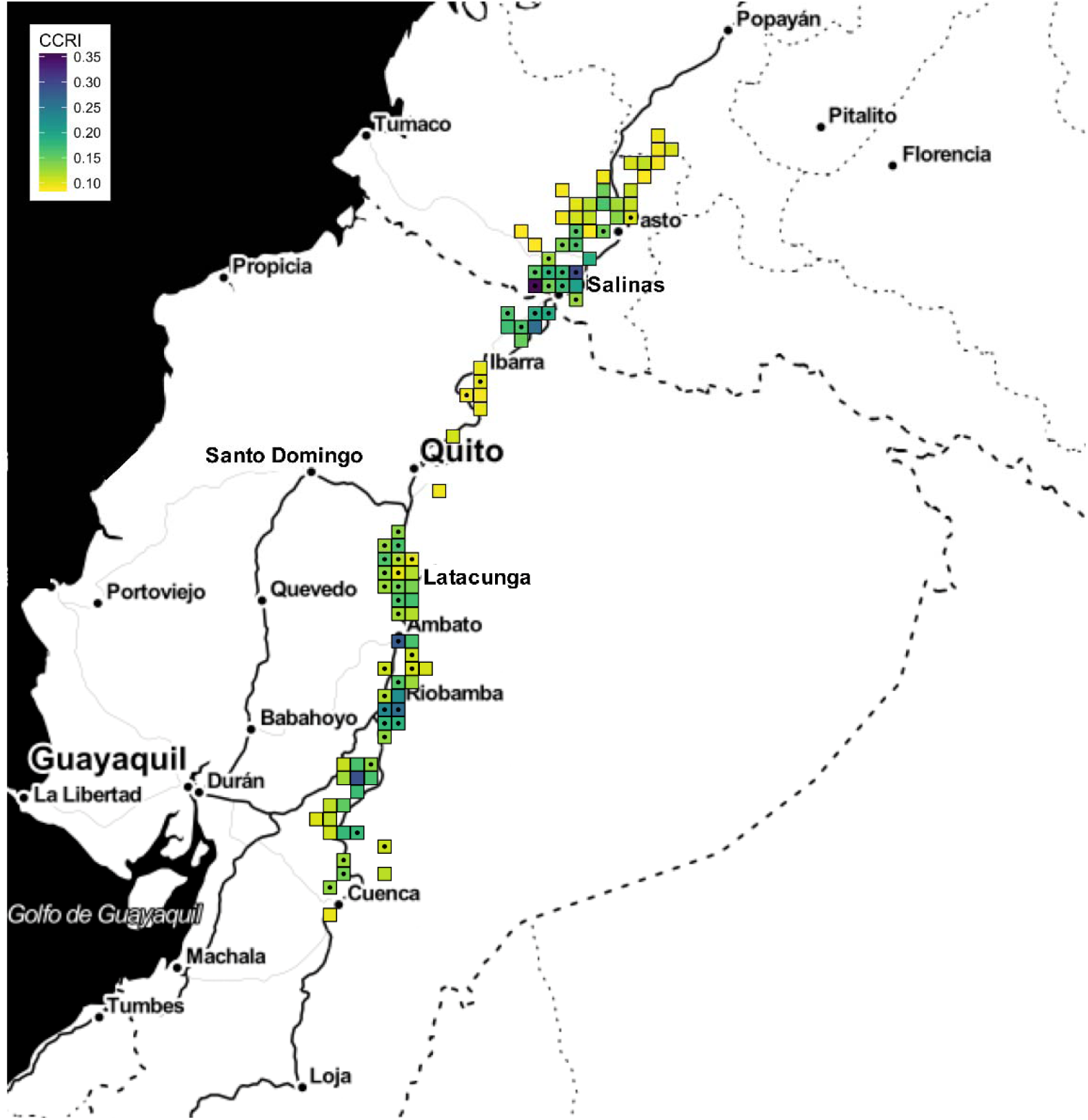
Ecuador and southern Colombia, with potato production indicated based on SPAM estimates. An altitude of 2895 m.a.s.l. was identified as a cut-off for management performance for positive selection of plants for on-farm seed saving. Pixels above 2895 m elevation (51% of the pixels) are indicated with a dot, where pixels are included if the harvested area estimate is greater than 200 ha. The graticules are 1-degree squares. Higher values of the potato cropland connectivity risk index estimated for Ecuador and southern Colombia are indicated by darker colors, indicating likely more important roles in potato epidemics. Targeting sites for farmer training in positive selection, might be based on the combination of being above the altitude cut-off for positive selection performance, and being in high cropland connectivity locations such that improved management would have the potential to positively influence other regions.

#### Technology adoption rates for three counties in Kenya

The county with the highest adoption rate for positive selection, Nakuru (Table 1), was intermediate in terms of the cropland connectivity index (Fig. 6). The cropland connectivity index was high for multiple locations in and near Nyandarua county.

**Figure 6.**
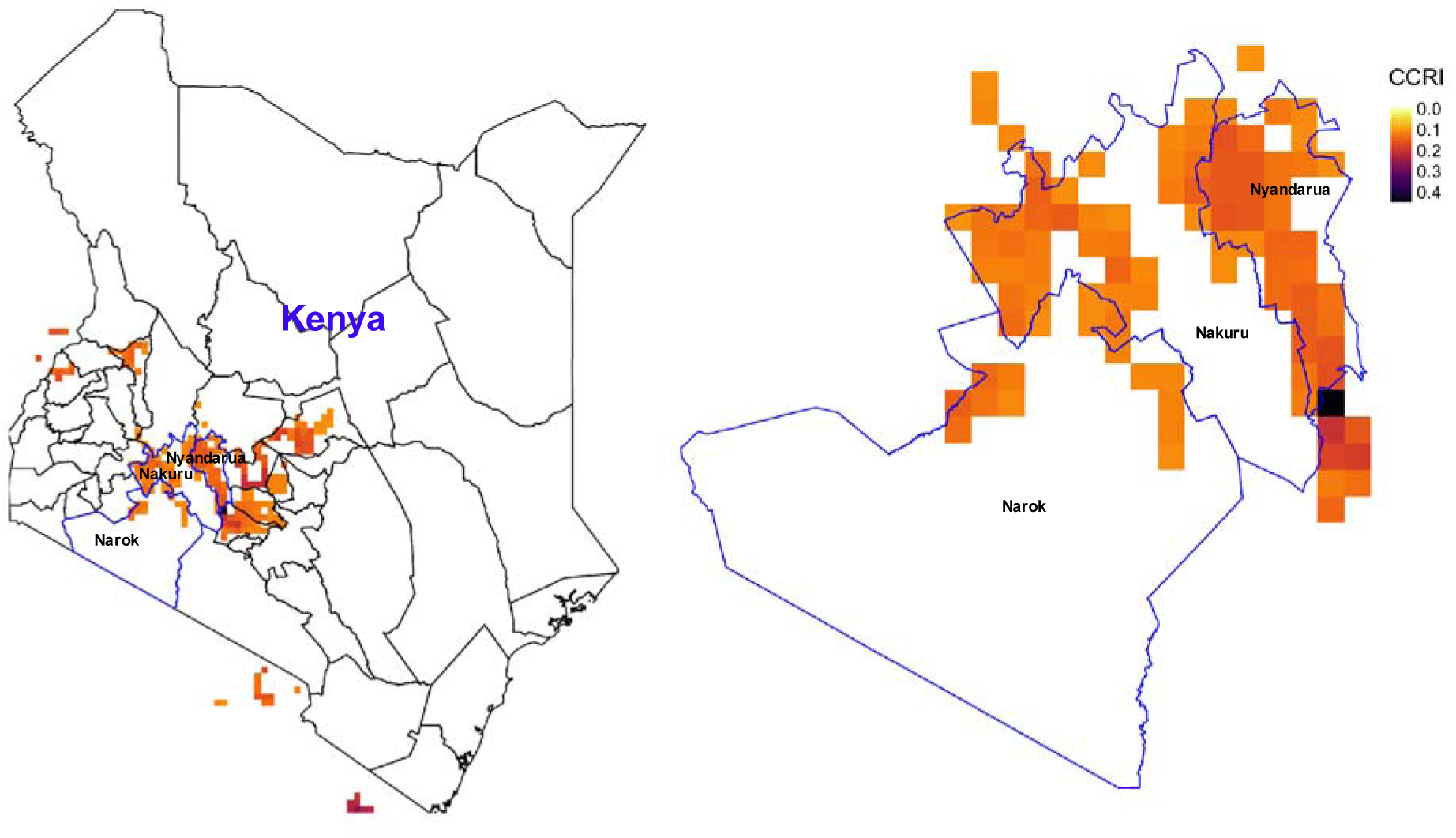
Cropland connectivity in the area of three counties in Kenya, where darker shading indicates a higher cropland connectivity risk index. Cropland connectivity is a measure of the likely importance of a pixel for epidemic spread through potato production. When the three counties indicated were studied to evaluate adoption rates for positive selection of plants for on-farm seed saving, Nakuru county was reported to have over twice the adoption rate. Targeting for training in positive selection methods could take into account the higher adoption rate in Nakuru and the higher cropland connectivity in Nyandarua.

### 5) Evaluate the value of information for management intervention or policy

#### VOI for targeted implementation of positive selection (yield as response, Bayesian networks to identify predictors)

For equivalent farm sizes above and below 2895 m (representing 49% of the area), we estimated the benefit of training under uninformed (random) site selection by using the weighted mean of the benefit above and below 2895 m.a.s.l. (representing 51% of the cultivated area), which is 6.5 t/ha (0.51 * 7.7 + 0.49 * 3.2 = 6.5).

The estimated benefit under informed site selection, selecting locations above 2895 m, is 7.7 t/ha – a difference of 1.2 t/ha from random site selection. Under misinformed site selection, if the assumption was that positive selection provides more benefits at low altitude (perhaps due to greater pathogen load), then the benefit is 3.2 tons per ha, 4.5 tons per ha less than the optimal allocation, and 3.3 tons per ha less than the uninformed (random) site selection option.

#### VOI for targeted implementation of positive selection (yield as response, recursive partitioning to identify predictors)

Assuming that positive selection training targeted farmers randomly with respect to the observed frequency of the categories, the weighted mean benefit of positive selection would be 8.7 tons per ha (Fig. 3). Preferentially targeting sites at high altitude (but sampling randomly with respect to seed age and cultivar) provides higher benefits (9 tons/ha), otherwise targeting low altitude sites provides lower returns at (8.3 tons/ha). Targeting farmers who plant farm-saved seed, three years since purchase of certified seed, provides little benefit: 8.8 tons/ha compared to random targeting of farmers (under the scenario where use of certified seed is rare at 2%). Targeting the 2% that do use new seed provides a benefit of 3.1 tons/ha, although these more successful farmers may not need interventions. By far the greatest benefit is provided by targeting farmers that grow INIAP-Fripapa (benefit of 10.9 tons/ha) as opposed to Superchola (benefit of 7.6 tons/ha).

#### Regional differences in adoption of training recommendations in Kenya

In the example of Kenyan potato seed technology adoption, targeted selection of high adoption rate areas for training (where the per-farmer benefit was $72) would increase the return by $28 per farmer trained (Table 1). Realized benefits would vary depending on the farm size.

## DISCUSSION

The examples given here show how management performance mapping can be used to target sites for project interventions. We illustrate how identifying locations where positive selection of on-farm saved seed has the highest performance for increasing yield (Andes) or the highest adoption rates (Kenya) can provide substantial regional benefits. While the particular examples here would require additional information to advance to the field with confidence (steps 6 through 8 in Fig. 1), they illustrate how a management performance mapping framework can be implemented An NGO or government extension agency with limited resources could use such an approach to better target rural development interventions. We compared uninformed and informed allocation of resources, for scenarios where the management performance models are correct (i.e., scenarios where the data perfectly represent the region of interest), to assess the value of the information used for targeting interventions. In a simple scenario, using Bayesian network analysis to identify altitude as a management performance predictor, we found that the benefit of positive seed selection was highest (an increase of 4.5 tons per ha) at high altitudes, and uninformed allocation of farmer training would provide a net benefit of 1.2 tons per ha less than targeted training. Incorrectly assuming that better outcomes for positive selection would be obtained at lower altitudes, perhaps because aphid vector abundances were thought to be higher, would have produced 3.3 and 4.5 tons per ha less for random site selection and optimal allocation, respectively, for this scenario (Bertschinger et al. 2017).

Along with the magnitude of management effects on yield, adoption rates are also key to successful interventions (Parsa et al. 2014). Based on the data about the benefits of adoption rates of positive selection of seed in Kenya, we found that unless adoption rates were higher than 24%, the first-year benefit per household would not exceed the $38 per farmer cost of training (although, presumably, the benefits would continue to accrue in subsequent years). Also, random allocation of training effort would only yield a $44 benefit (over the cost of the training) per household. Gildemacher et al. (2017) also point out that adoption rates were lower in drought years, suggesting that prediction of adoption rates could be difficult if based on regional patterns in a single year. Observed adoption rates may vary in predictable ways based on disease incidence in the current or previous season, in-season weather conditions, language spoken, literacy, cultural differences between trainer and trainee, wealth or other factors. When these relationships are understood and spatial data are available for key predictor variables for adoption, these variables could form a part of selection criteria for farmer training initiatives (and the approach to the training could be altered to improve adoption rates).

Our example decision, deciding where to implement training for improved disease management, represents a class of decisions where there is confidence that the activity will provide a benefit. Management performance mapping is applied to guide implementation to locations where there is some evidence that the benefit will be greater than in other locations. For this class of decisions, the risk is often low that limited data is “worse than no data at all”. In the management performance mapping context, the null hypothesis is often that the benefit of implementation will be the same in all locations. In evaluating where there is evidence to reject this hypothesis, there is not a strong motivation to particularly avoid false positives or Type I error (rejecting a null hypothesis when the null hypothesis is true), because a false negative or Type II error (failing to reject a null hypothesis when the null hypothesis is false) is arguably just as bad. The main risk of “bad data” would be from data with a strong bias that would lead to misinformed decisions. The cost of “bad data” may also go up if the logistical costs (of transport, communications, etc.) of targeting locations incorrectly identified is higher than targeting locations at random or selecting locations based on convenience. There is the potential for these risks to be managed in real time during project implementation by incorporating distributed or “big” data sourced from farmer phone apps, rapid disease detection methods or citizen science initiatives (Nakato et al. 2016; Boykin et al. 2019). In our example data from Ecuador, our only estimate of uncertainty within a scenario was based on variability among individual plants, while a person making decisions about regional priorities would strongly prefer to have information about farm-to-farm variability within each scenario. One of the potential applications of VOI analysis is to determine whether collecting more or better data about management performance is justified (Ades et al. 2004), not just for the sake of more statistical power in general, but because the information improves farmer decision-making under a realistic range of conditions.

Two key factors for adoption of positive selection are market price and the varieties grown in a region, in terms of their rates of seed degeneration. The number of seasons over which positive selection is adopted is also an important factor helping to determine the return on investment in training. The study of adoption rates in Kenya was performed once and may be limited to the circumstances at the time of sampling. At the time of the training in positive selection, there was a shift in Nyandarua from the variety Tigoni to Shangi, so there might have been more interest in acquiring the new variety than in improving old seed stock (Okello et al. 2018; Kaguongo et al. 2008). Nyandarua also has an apparent role in spread of potato cyst nematode in the region (Mburu et al. 2018; Mwangi et al. 2015), along with bacterial wilt problems, which may have made positive selection less effective there. Potato farming has a longer history in Nyandarua. In Narok, potatoes are less important and conditions are less favorable, with the main potato variety grown being Dutch Robijn. In Nakuru, with the highest adoption rate, more varieties are grown and potato farming is more recent, in generally good growing conditions. These differing factors in the three counties, combined with changes over time such as the occurrence of droughts, can modify the likelihood of adoption of positive selection. The yield improvements from positive selection in Kenya, averaging 30% (Schulte-Geldermann et al. 2012), make it an attractive technology for development investments. In this system, there is the possibility of farmers actually improving seed quality over time, rather than simply slowing decline, and understanding this potential could also support decisions. Formulating a strategy for targeting positive selection in Kenya would be strengthened by new data about how the differences among these and other counties influence the current likelihood of technology adoption.

Combining data layers for evaluating optimal intervention strategies can provide more insight, along with potential challenges due to uncertainty and different spatial resolutions (Sutton and Armsworth 2014). Evaluating the risk of disease due to cropland connectivity (Xing et al. 2020) in combination with independent location characteristics can position the analysis in the larger context of disease management for the region. Cropland connectivity may change over the course of the year, as potato is present or absent. For example, in Kenya some parts of Nyandarua county, such as Njambini and Oljororok, have potato in the field the entire year, due to the availability of groundwater coming from the Aberdare range during the dry season. Three crops a year are very common, and likely affect pest and disease cycles. Consistently highly connected locations may be more important targets for achieving impacts on regional epidemic spread, although there is also the potential for highly connected locations experiencing high inoculum loads to respond poorly to some types of management. A broader systems analysis – for example, impact network analysis (Garrett et al. 2018) which integrates across management performance, socioeconomic or innovation networks (Fritsch and Kauffeld-Monz 2010; Leeuwis and Aarts 2011), and biophysical networks such as epidemic networks – can aid in identifying intervention locations that prioritize across multiple goals. For farmer decision making, flexible decision rules and reducing variability of risk may be priorities (Bert et al. 2006; Andrieu et al. 2015).

Crop and epidemic models may provide valuable data layers if they incorporate spatially mappable variables. Estimating the effects of management strategies, such as variety deployment, depends on understanding the yield potential, perhaps based on a combination of weather or climate data and data about regional management practices (van Wart, van Bussel, et al. 2013; Araya et al. 2010; Reynolds et al. 2018). Disease modeling may be used to evaluate the likely effects of management, such as addressing the problem of seed degeneration (Thomas-Sharma et al. 2017; Jones et al. 2010). Spatial epidemic components such as seed trade networks (McQuaid et al. 2017; Buddenhagen et al. 2017; Andersen et al. 2019) may also be valuable components of more refined management performance mapping.

Management performance mapping to identify target locations for interventions, and the VOI analysis therein, is potentially useful for many problems in agriculture or intervention ecology. There is a constellation of approaches that address related goals. Yield gap analyses that incorporate maps can address some of the same goals as management performance mapping (Schulthess et al. 2013; Silva et al. 2017; Lobell et al. 2015, 2009; van Ittersum et al. 2016; Grassini et al. 2015; van Bussel et al. 2015). For example, yield gap analysis attempts to identify the most important factors that influence yield, especially factors that are controllable. The focus of management performance mapping for intervention targeting, however, is on providing spatial information about the intervention impact of management options. Management performance maps would ideally incorporate and account for interacting human dimensions (e.g., learning, financial liquidity, capital, institutions; Arneth et al. 2014), as well.

The benefits from management performance mapping may be enhanced if maps and VOI calculations are updated with new information sources over time in an adaptive management scheme (Shea et al. 2014; Bennett et al. 2018). Empirical data can be supplemented with models, expert opinion, and local knowledge (Petsakos et al. 2018; Tulloch et al. 2014) to understand changes in factors such as pesticide resistance, new varieties, and new management such as irrigation. Projects may expand due to new stakeholder priorities and new situations on the ground. As new pests and pathogens enter a region (Bebber et al. 2014), they will likely necessitate alterations in current best management practices. For example, in Ecuador potato purple top disease has become a major problem (Caicedo et al. 2015; Castillo-Carrillo et al. 2018) since the Ecuadorian experiments reported here were performed. ‘Candidatus Liberibacter solanacearum’, associated with zebra chip disease, and its vector the tomato potato psyllid, *Bactericera cockerelli*, have also been reported in Ecuador (Caicedo et al. 2020; Castillo-Carrillo et al. 2019). New strategies for potato best management practices in Ecuador will need to address purple top and the risk of zebra chip, including uncertainty about causal agents. In other scenarios, multiple outcomes may be important, such as a combination of benefits and environmental costs of management (Laurance et al. 2014), pesticide effects on non-target species in disease management, or conservation management focusing on both biodiversity hotspots and locations with keystone species (Smith et al. 2007). Our examples addressed management performance mapping with performance defined in terms of the <u>mean</u> performance observed. Other potential criteria for selecting regions for investment might emphasize different priorities (Table 2). For example, effective altruism concepts can be used to target stakeholders to maximize research benefit (Garrett et al. 2020).

**Table 2.** Potential criteria for identifying priority sites for interventions (such as training farmers to use positive selection for improved seed health).

| **Criterion** | **Rationale** |
| --- | --- |
| Regions where expected absolute benefit is greatest | Greatest benefit to regional food production |
| Regions where expected proportional gain is greatest | Greatest benefit to regional farmers |
| Regions where outcomes before intervention are lowest | Benefit to regions in greatest need |
| Regions where outcomes before intervention are highest | Benefit to regions currently best adapted for production |
| ***Incorporating measures of uncertainty at the farm level*** |  |
| Regions where the 5<sup>th</sup> percentile benefit is greatest | Consistent benefit across farmers |
| Regions where the trimmed mean is greatest | Greatest benefit for typical farmers |

In summary, management performance mapping provides a process to extrapolate from available data to make evidence-based decisions about where to invest in disease and crop management or training initiatives. Scenario analyses to support decision making (Wiebe et al. 2018) can build on the framework developed here.

## ACKNOWLEDGMENTS

This research was undertaken as part of, and funded by, the CGIAR Research Program on Roots, Tubers and Bananas (RTB) and supported by CGIAR Trust Fund contributors https://www.cgiar.org/funders/. We also appreciate support by the CGIAR Research Program on Climate Change and Food Security (CCAFS), Bill and Melinda Gates Foundation grant OPP1080975, USDA NIFA grant 2015-51181-24257, USDA APHIS grant 11-8453-1483-CA, the USAID Feed the Future Haiti Appui à la Recherche et au Développement Agricole (AREA) project grant AID-OAA-A-15-00039, US NSF Grant EF-0525712 as part of the joint NSF-NIH Ecology of Infectious Disease program, US NSF Grant DEB-0516046, and the University of Florida.

